# Dissociation of solid tumour tissues with cold active protease for single-cell RNA-seq minimizes conserved collagenase-associated stress responses

**DOI:** 10.1101/683227

**Authors:** Ciara H O’Flanagan, Kieran R Campbell, Allen W Zhang, Farhia Kabeer, Jamie LP Lim, Justina Biele, Peter Eirew, Daniel Lai, Andrew McPherson, Esther Kong, Cherie Bates, Kelly Borkowski, Matt Wiens, James Hopkins, Brittany Hewitson, Nicholas Ceglia, Richard Moore, Andy J Mungall, Jessica N McAlpine, The CRUK IMAXT Grand Challenge Team, Sohrab P Shah, Samuel Aparicio

## Abstract

**Background:** Single-cell RNA sequencing (scRNAseq) is a powerful tool for studying complex biological systems, such as tumour heterogeneity and tissue microenvironments. However, the sources of technical and biological variation in primary solid tumour tissues and patient-derived mouse xenografts for scRNAseq, are not well understood. Here, we used low temperature (6°C) protease and collagenase (37°C) to identify the transcriptional signatures associated with tissue dissociation across a diverse scRNAseq dataset comprising 128,481 cells from patient cancer tissues, patient-derived breast cancer xenografts and cancer cell lines.

**Results:** We observe substantial variation in standard quality control (QC) metrics of cell viability across conditions and tissues. From FACS sorted populations gated for cell viability, we identify a sub-population of dead cells that would pass standard data filtering practices, and quantify the extent to which their transcriptomes differ from live cells. We identify a further subpopulation of transcriptomically “dying” cells that exhibit up-regulation of MHC class I transcripts, in contrast with live and fully dead cells. From the contrast between tissue protease dissociation at 37°C or 6°C, we observe that collagenase digestion results in a stress response. We derive a core gene set of 512 heat shock and stress response genes, including *FOS* and *JUN*, induced by collagenase (37°C), which are minimized by dissociation with a cold active protease (6°C). While induction of these genes was highly conserved across all cell types, cell type-specific responses to collagenase digestion were observed in patient tissues. We observe that the yield of cancer and non-cancer cell types varies between tissues and dissociation methods.

**Conclusions:** The method and conditions of tumour dissociation influence cell yield and transcriptome state and are both tissue and cell type dependent. Interpretation of stress pathway expression differences in cancer single cell studies, including components of surface immune recognition such as MHC class I, may be especially confounded. We define a core set of 512 genes that can assist with identification of such effects in dissociated scRNA-seq experiments.

## Introduction

Recent advancements in sequencing technologies have allowed for RNA sequencing at single cell resolution, which can be used to interrogate features of tumour tissues that may not not be resolved by bulk sequencing, such as intratumoural heterogeneity, microenvironmental architecture, clonal dynamics and the mapping of known and de novo cell types. Due to the sensitivity of single cell RNA sequencing (scRNAseq), small changes in gene expression can dramatically influence the interpretation of biological data. scRNAseq data is also subject to technical and biological noise [1, 2]. The inherent nature of the transcriptome is transient and dynamic, reflecting the ability of cells to quickly respond to their environment. In addition, the transcriptional behaviour of single cells can deviate profoundly from the population as a whole, and gene expression pulse patterns have been shown to contribute significant noise levels to scRNAseq data [3]. Inherent variations in tissue composition, cell quality and cell-cell variability can also make it difficult to confidently interpret scRNAseq data. While current technologies attempt to mitigate noise from amplification during library construction by the incorporation of unique molecular identifiers (UMIs) during cDNA synthesis [4], this does not address changes to the transcriptome prior to reverse transcription. High-quality scRNAseq data requires highly viable single cell suspensions with minimal extracellular components, such as debris. Standard sample preparation methods for solid tissues require enzymatic and mechanical dissociation, and depending on the tissue origin, density, disease state, elastin or collagen content, may require long enzymatic digestion and/or vigorous mechanical disruption. Transcriptional machinery remains active at 37°C, and extended incubation at high temperatures may introduce gene expression artifacts, independent of the biology at the time of harvest. Moreover, extended incubation at higher temperatures in the absence of nutrients or anchorage, or harsh dissociation may induce apoptosis or anoikis, polluting the viable cell population or generating low quality suspensions [5]. Therefore, it is imperative to characterize the inherent variation and potential effects of cell isolation methods on the transcriptomic profiles of tissues. Recently it has been shown that a serine protease (subtilisin A) isolated from a Himalayan glacier-resident bacterium, *Bacillus Lichenformis*, is suitable for dissociation of non-malignant renal tissues at 4-6°C, and can reduce scRNAseq artifacts in these tissues, including reducing global and single cell gene expression changes [6].

Given the heterogeneous nature of tumour tissue [7–9], and the potential application of scRNAseq in studying the complex biology of cancer including the tumour microenvironment [10], tumour heterogeneity [9] and drug response [11], we sought to determine the effects of of enzymatic dissociation and temperature on gene expression artifacts in tumour tissues and cell lines. Here, using a diverse scRNAseq dataset of 40 samples and 128,481 cells comprising patient cancer tissues, patient-derived breast cancer xenografts (PDXs) and cancer cell lines, we highlight the inherent variation in scRNAseq quality control metrics across samples and constituent cell types in patient tumour samples. We identify a subpopulation of dead cells that would not be removed through standard data filtering practices and quantify the extent to which their transcriptomes differ from live sorted cells. We identify a further subpopulation that represents transcriptomically dying cells, expressing increased major histocompatibility complex (MHC)-class I genes. We identify a core geneset of immediate, heat shock and stress response genes associated with collagenase dissociation, highly conserved across cell and tissue types, which are minimized by dissociation at cold temperature. These findings may significantly affect biological interpretation of scRNAseq data and should taken into careful consideration when analyzing single cell experiments.

## Results

### Single cell RNA-sequencing of 128,481 cells

To uncover transcriptional variation and responses to dissociation method, we generated scRNAseq data for 128,481 single cells across a range of substrates, cancer types, dissociation temperatures, and tissue states (**Figure 1**), using the 10X Genomics Chromium v3 platform [12]. scRNAseq was performed on cells from patient samples, PDXs and cell lines across ovarian, lymphoid cell and breast cancers, including fresh and viably frozen samples dissociated at 37°C or 6°C and cells incubated at 6°C, 24°C, 37°C, or 42°C (Figure 1).

**Figure 1:**
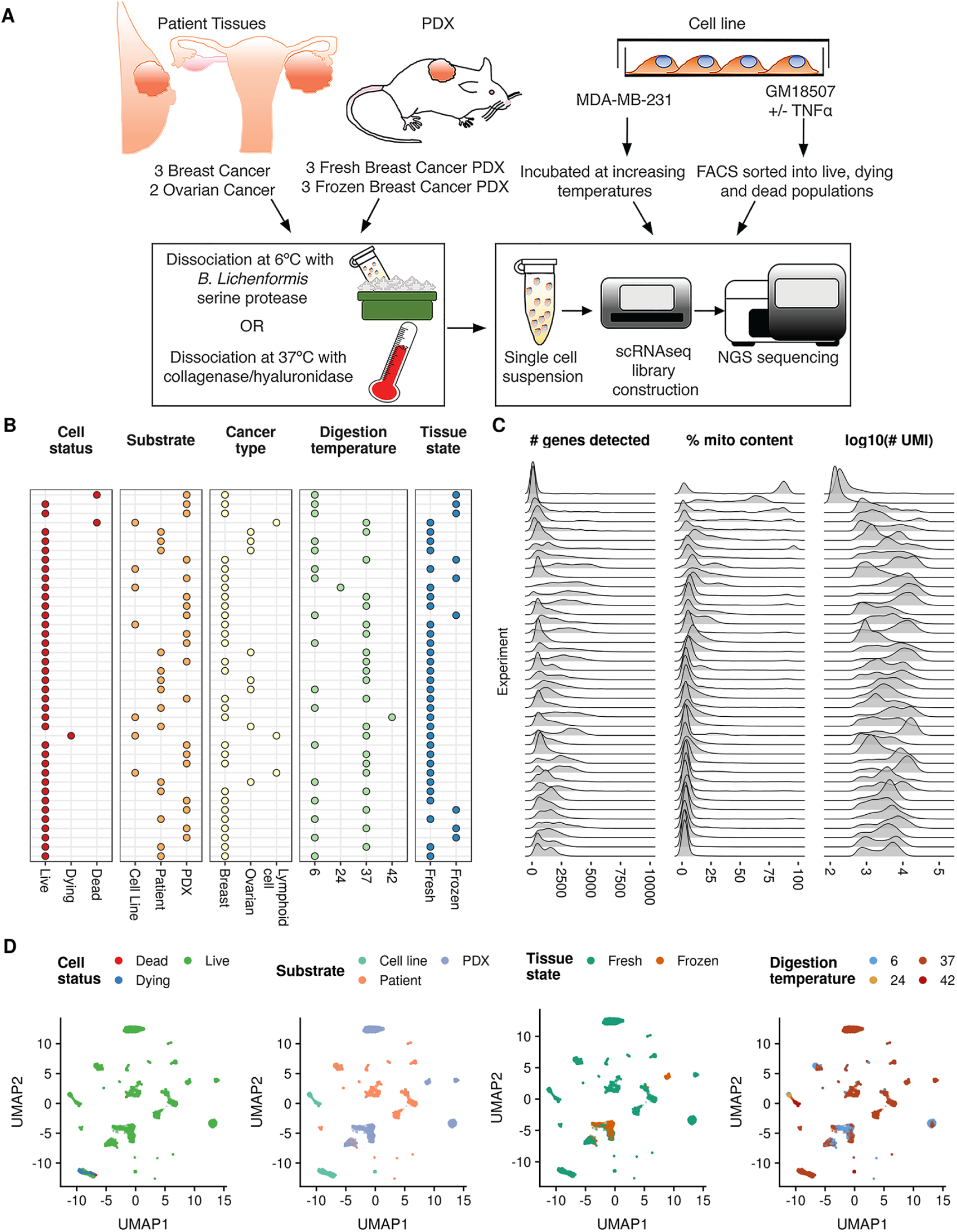
Overview of 40 single-cell experiments generated in this study. **A** Schematic showing the various substrates used to generate the 40 single-cell experiments in this dataset. **B** Descriptions of the cell status, substrate, cancer type, dissociation temperature and tissue state of each sample in the dataset. **C** Substantial variability in 3 key QC metrics (number of genes detected, percentage of counts mapping to the mitochondrial genome, number of UMIs sequenced) across all experiments. **D** Embedding of all 40 single-cell experiments to a low-dimensional projection with uniform manifold approximation and projection [13].

We began by examining a set of commonly used quality control (QC) metrics across all 40 sequencing experiments (**Figure 1C**), including total number of genes detected, percentage of transcripts mapping to the mitochondrial genome, and total number of UMIs sequenced. We observed significant variation across these metrics, in particular bi- and tri-modal distributions of mitochondrial gene percentages across this varied sample set. This variable mitchondrial gene content was also observed in publicly available datasets from 10x Genomics (Figure S1).

Conscious of the possibility of murine stromal cell contamination in PDX samples, we classified cells as mouse or human based on alignment metrics. Of the 55,332 PDX cells sequenced, 1,208 were reliably identified as mouse cells, with large inter-sample variation (Figure S2). We found 372 cells across primary tumour and cell line samples were mis-identified as murine compared to 69,608 cells identified as human, suggesting this approach to detecting murine contamination has a modest false positive rate of 0.5%. As expected, murine cells scored consistently lower across a range of standard QC metrics (% counts mitochondrial, total genes detected, total UMIs detected) when aligned to the human genome (Figure S3).

### Transcriptomic landscape of live, dead, and dying cells

Given the bi- and tri-modal distributions of mitochondrial gene count percentages apparent in the 40 experiments and previous studies’ assertions that high mitochondrial gene content is indicative of dead and dying cells [14, 15], we next sought to determine the contribution of dead and dying cells to the variation observed in QC metrics in **Figure 1**. In order to induce classical cell death pathways, we used TNFα[16, 17] to treat the non tumourigenic, lymphoblastoid cell line GM18507 and FACS sorted cells into dead or and dying fractions based on PI/Annexin V positivity (**Figure 2A**), as well as a live, untreated fraction. Notably, cell yield from scRNAseq data was highly dependent on the cell status, with 8,597 live cells recovered but only 1,280 and 885 dead and dying respectively compared to targeted numbers of 3,000 cells.

**Figure 2:**
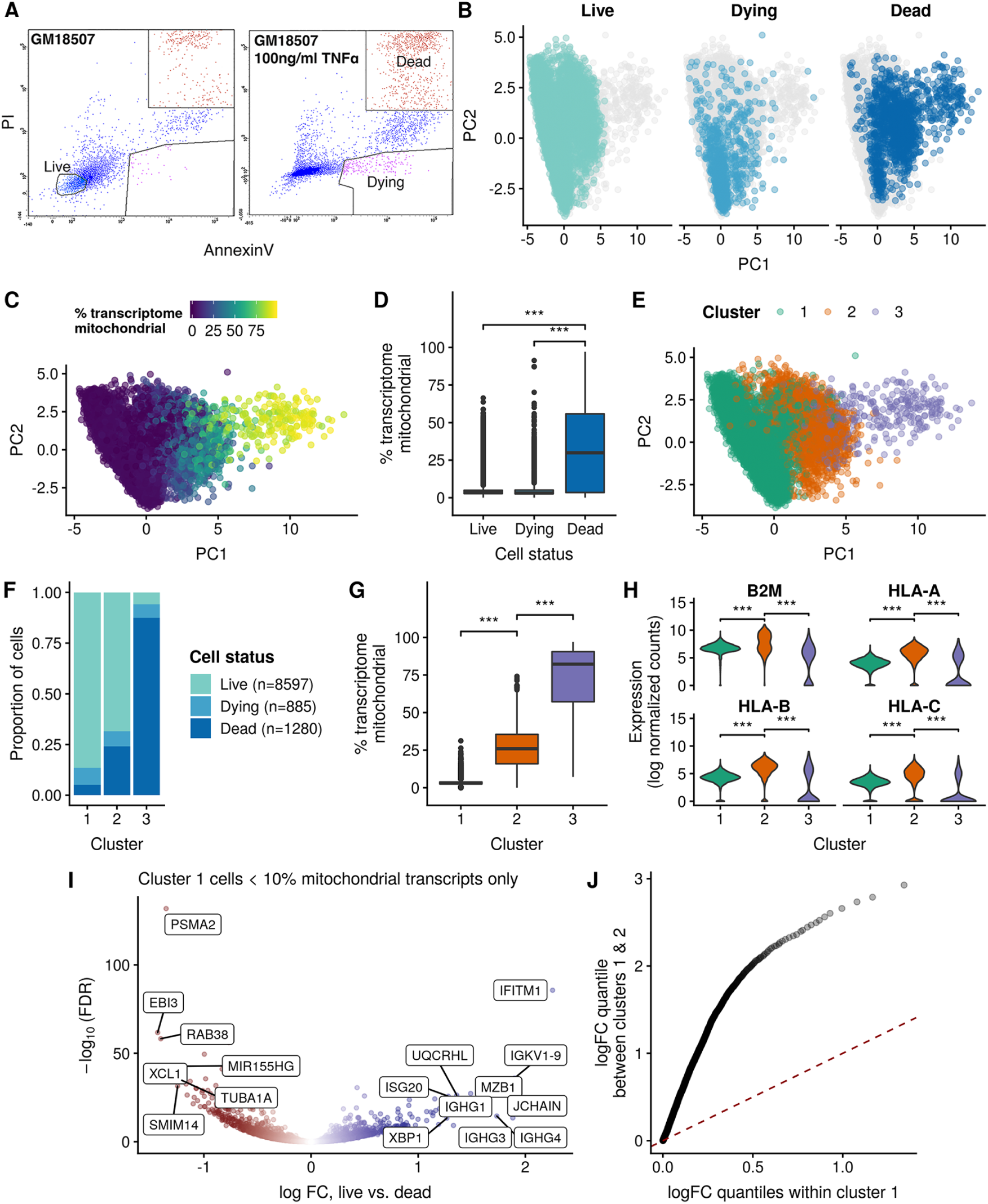
Transcriptomic landscape of live, dead, and dying cells. **A** FACS analysis showing gating strategy for untreated, live cells (PI-/AnnexinV-) or TNFα-treated dying cells (PI-/AnnexinV+) and dead cells ((PI+/AnnexinV+). **B** PCA projection of the 3 cell conditions showing approximate segregation of cell status along the first principal component (PC1), with live and dying cells enriched at lower PC1 values and dead cells enriched at higher values. **C** PCA projection coloured by the percentage mitochondrial genes (“% transcriptome mitochondrial”) shows significant increase along the PC1. **D** Dead cells exhibit significantly higher % of the transcriptome as mitochondrial compared to both live and dying cells. **E** Unsupervised clustering of the gene expression profiles clusters the cells into 3 groups, approximately tracking both PC1 of the data and the % transcriptome mitochondrial. **F** The composition of each cluster demonstrates that cluster 1 is primarily composed of live cells, cluster 2 a mix of live, dying, and dead cells, while cluster 3 is composed mainly of dead cells. **G** The % transcripts mitochondrial is significantly different between the three clusters, with a step increase in proportion moving from cluster 1 to 2 and 2 to 3. **H** Cluster 2 significantly up-regulates the MHC class I gene set, suggesting it represents stressed or pre-apoptotic cells. **I** Differential expression analysis of transcriptomically “healthy” cells within cluster 1 reveals residual differences between cells sorted as live and dead. **J** The distribution of absolute effect sizes (log fold change) of live vs. dead cells within cluster 1 (x-axis) compared to between clusters 1 and 2 (y-axis) demonstrates the residual effect on the transcriptome of being live/dead sorted is small compared to the inter-cluster expression variance.

A principal components analysis (PCA) following mutual nearest neighbours (MNN) correction [18] demonstrated the cells approximately segregating along the first principal component (PC1) by cell status (**Figure 2B**), albeit with high levels of heterogeneity in overlap. Indeed, PC1 closely tracked the mitochondrial gene content of the cells (Figure 2C), being significantly higher in dead cells (median 29.9%) compared to both dying cells (median 3.13%, *p* =1.17e-126) and live cells (median 3.4%, *p* = 4.65e-153) as shown in **Figure 2D**. This observation justifies the practice of excluding cells with very high mitochondrial gene content as being likely dead cells.

Having observed that the transcriptomes of the different cell conditions are not entirely distinct, we sought to discover the extent of mixing between transcriptomic states and whether live and dead cells that appear transcriptomically “healthy” (ie would ordinarily pass QC) are distinguishable. Using hierarchical clustering (methods) we clustered the cells into 3 groups that approximately track PC1 (**Figure 2E**). Interestingly, these three groups show variable composition in terms of cell states, with cluster 1 being comprised mainly of live cells (86% live, 8.5% dying, 5.1% dead), cluster 2 containing an increased proportion of dying and dead cells (68% live, 7.5% dying, 24% dead), and cluster 3 comprised mainly of dead cells (5.9% live, 6.7% dying, 87% dead). Furthermore, we observed a step change increase in mitochondrial gene content between clusters (**Figure 2G**), with cluster 1 having the lowest (median 3.13%), followed by cluster 2 having a significant increase (median 26%, *p* =0) and cluster 3 having a significant increase beyond that (median 82.2%, *p* =2.35e-149). Differential expression analysis between these clusters revealed a significant up-regulation in stress-associated pathways such as MHC class I (**Figure 2H**) in cluster 2 compared to clusters 1 & 3. MHC class I genes are involved in antigen presentation to T cells, but are also expressed in many cell types and are induced in response to stress stimuli and contain heat shock-inducible elements [19].

Together, these results suggest a model whereby cluster 1 represents transcriptomically “healthy” cells, cluster 2 represents transcriptomically stressed cells that upregulate stress pathways and have increased mitochondrial gene content (due to either an increasingly permeable membrane causing loss of cytoplasmic mRNA or increased metabolic demands), and cluster 3 represents transcriptomically dead cells whereby the membrane has burst leaving majority mitochondrial transcripts. Importantly, cells that are FACS sorted as either live, dying, or dead, are present in all three clusters, highlighting that the transcriptomic state of the cell is not necessarily the same as the surface marker state (though the two are correlated). Such concepts are reminiscent of “pseudotime” in single-cell developmental biology, whereby developmentally ordering cells transcriptomically can lead to early or late cells being placed at variable positions along the pseudotime trajectory [20, 21]. Indeed, PC1 from **Figure 2A** approximates a pseudotime trajectory through the data, that tracks transcriptomically healthy cells to transcriptomically dead cells with increasing PC1 values.

Finally, we sought to determine if a sorted dead cell that appears transcriptomically healthy remains distinguishable from a sorted live cell in the transcriptomically healthy group. Using only cells in cluster 1, we further subsetted them to pass a strict set of QC filters (at least 10^3^ total genes detectable, % mitochondrial content between 1 and 10) and performed a differential expression analysis between cells sorted as live and dead in this group. Of the 10537 genes retained for analysis, 2130 (20.2%) were found to be differentially expressed (**Figure 2I**), including downregulation of *IFITM1* in dead cells. To compare this type of variation to the inter-cluster transcriptomic variation, we performed a second differential expression analysis between clusters 1 and 2, finding 8835 of 10933 (80.8%) genes significantly differentially expressed. Furthermore, the effect sizes were significantly larger for the inter-cluster comparison than the within-cluster 1 live-dead comparison as demonstrated by the quantile-quantile plot of absolute effect sizes in figure Figure 2J. Together, these results suggest that though there are gene expression differences between dead and live sorted cells within cluster 1, the magnitude of expression variation is small compared to transcriptomically stressed clusters.

### Dissociation with collagenase at 37°C induces a distinct stress response in single-cell transcriptomes

To uncover the effect of digestion temperature on the transcriptome, we performed a differential expression analysis on the 23,731 cells found by combining all experiments measured in a PDX or cell line at either 6°C or 37°C. We removed any samples corresponding to primary tumours as we discovered that yield of constituent cell types was affected by digestion temperature (Figure S6), which would confound our differential expression results. After retaining genes with at least 10 counts across all cells, we performed differential expression analysis with edgeR [22], while controlling for the sample-of-origin.

We found that of the 19,464 genes retained for analysis, 11,975 (62%) were differentially expressed at a Benjamini-Hochberg corrected false discovery rate (FDR) of 5%. We defined a core set of genes meaningfully perturbed by digestion temperature as those significantly differentially expressed as above, but with an absolute log fold change of at least 1.5. Therefore, for a gene to be included under these criteria it must be differentially expressed and its abundance increased or decreased by at least 50% by digestion temperature. This produced a core gene set of 512 genes, of which 507 were upregulated at 37°C and the remaining 5 downregulated. This gene set includes multiple canonical stress-related genes such as *FOS*, *FOSB*, *ATF3* and heat shock proteins (HSPs) (Figure 3A), expression of which have shown to be induced by collagenase dissociation in a subset of muscle cells [23]. A UMAP embedding of the cells coloured by dissociation temperature and the expression of several key genes (*FOS*, *JUNB*, *NR4A1*, Figure 3B) further demonstrates the digestion temperature-specific induction of the expression of these genes.

**Figure 3:**
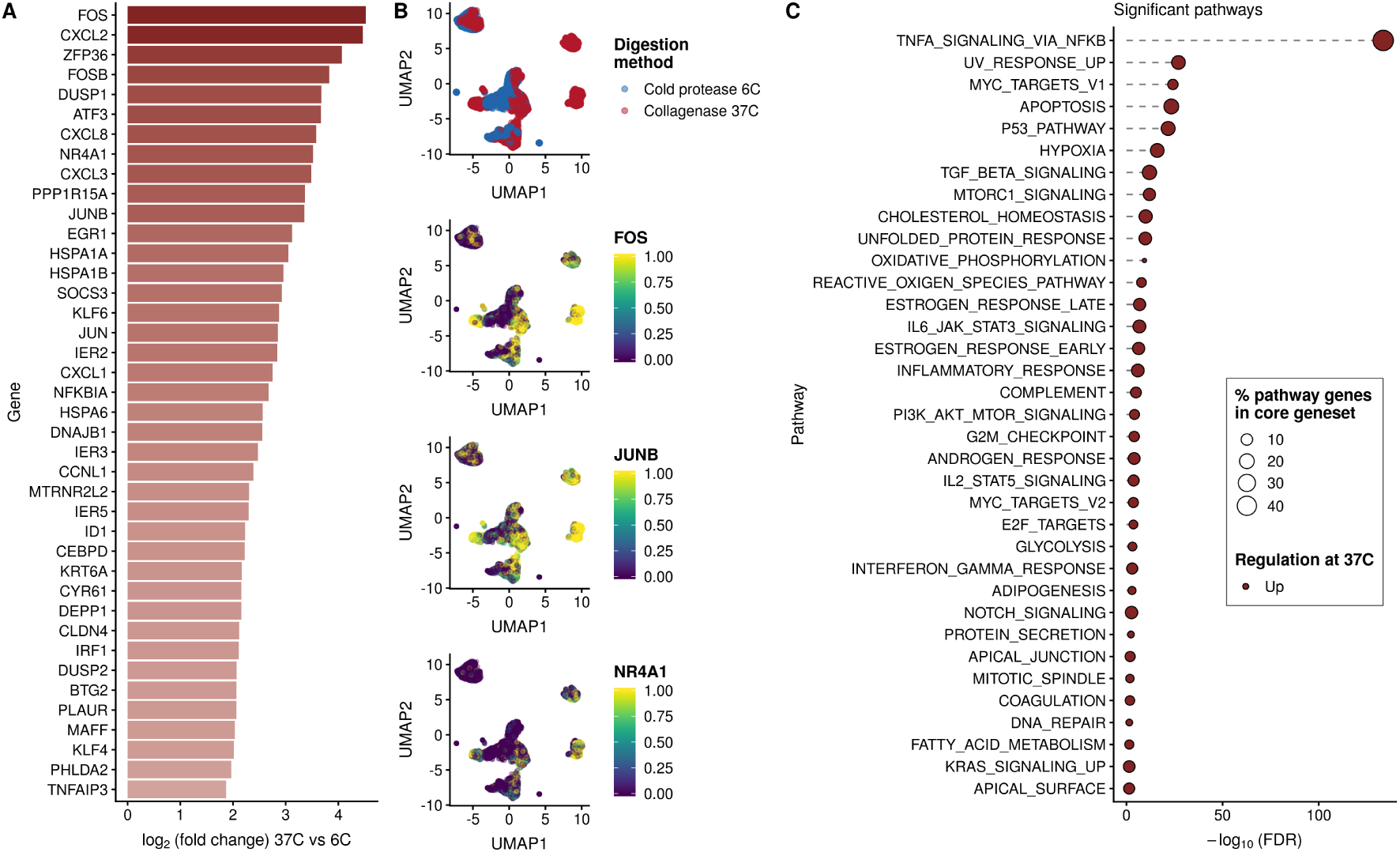
Dissociation with collagenase at 37°C induces a distinct stress response in 23,731 cells from PDX samples that is minimized by dissociation at 6°C. **A** The top 40 genes (by log fold-change) from the 11,975 identified as significantly differentially expressed between cells digested at 6°C and 37°C. **B** UMAP plots of ncellspdxde cells coloured by digestion temperature (top) then by normalized expression of 3 key stress response genes (*FOS*, *JUNB*, *NR4A1*) demonstrates a distinct concordance between temperature and induction of the stress gene signature. Expression values are log normalized counts (**methods**) winsorized to [0, 2) then scaled to [0, 1). **C** Pathway analysis of differentially expressed genes with the MSigDB hallmark gene sets highlights induction of genes involved in NF-*κ*B signalling at 37°C digestion with 46.5% of 200 genes annotated in the pathway being found in the 512 core gene set.

Noting the large number of HSP proteins significantly upregulated at the 37°C collagenase digestion, we examined their expression in the MDA-MB-231 samples incubated at different temperatures (6°C, 24°C, 37°C, 42°C). Upregulation of the HSP genes in the 512 core geneset typically follows a step increase between 37°C and 42°C incubation rather than a gradual increase with increasing temperature (Figure S4), implying their induction at 37°C collagenase digestion is due to a different mechanism than the digestion temperature alone, consistent with previous results [23].

We subsequently performed a pathway enrichment analysis on the differential expression results, searching for enrichments in given hallmark pathways [24] (Figure 3C). Of particular note was TnF signalling via NF-*κ*B of which 46.5% of annotated pathway genes were included in the core set of 512 genes (Figure S5). Further enrichment of stress-associated pathways including Hypoxia, Apoptosis, and Inflammatory response is further indicative of collagenase dissociation at 37°C as inducing a stress response on the transcriptomes of single cells.

### Conserved stress response to collagenase dissociation method in breast and ovarian patient tissues

Having derived a core geneset of stress and heat shock genes induced in PDX samples during dissociation with collagenase, we next examined the effect of dissociation method on recovery and transcriptomes of constituent cells of the tumour microenvironment in breast and ovarian patient samples. Histology and FACS analysis revealed a complex and variable tumour microenvironment (Figure 4A-B). Dissociation of ovarian cancer sample with cold protease yielded enhanced capture of lymphocytes including T cells, cytotoxic T cells, and NK cells (**Figure 4B**, Figure S6). We generated scRNAseq data of 2 high grade serous ovarian (HGSC) and 3 breast cancer samples (table S1) dissociated using collagenase at 37°C or cold protease at 6°C as described above. Total cell yield was highly variable, ranging from 282 to 9,640 cells across samples. Cells were subsequently assigned to a range of tumour microenvironment cell types using CellAssign [25], assuming a set of common marker genes for cell types (table S2, table S3). A UMAP project of the data (**Figure 4C**) demonstrates the broad range of cell types identified from the scRNA-seq data, including epithelial cells, structural cell types such as endothelial and myofibroblast cells, and an array of immune cell type such as B cells, T cells, Monocyte/Macrophage populations and plasma cells, consistent with FACS analysis (**Figure 4B**). While enhanced capture of certain lymphocyte populations was apparent in ovarian samples dissociated at 6°C, overall microenvironment composition was highly variable both between patients, reflected in histological analysis (**Figure 4A**), and dissociation protocols (Figure S6), no consistent loss or gain of cell types was observed between conditions in all samples.

**Figure 4:**
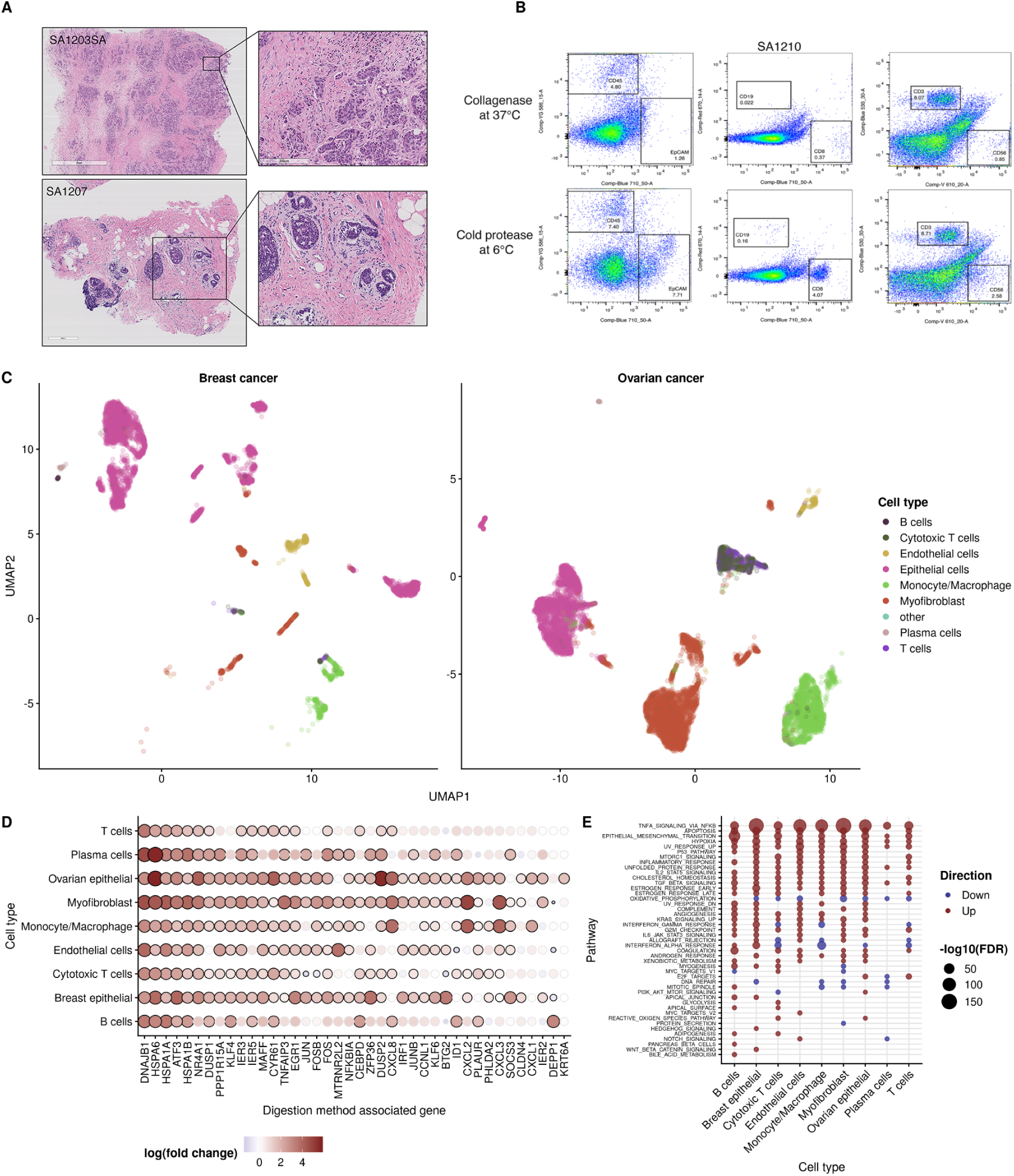
Conserved stress response to collagenase dissociation method in breast and ovarian patient tissues. **A** Histology of ovarian cancer patient sample highlighting the architecture of the tumour microenvironment. **B** FACS analysis of ovarian tumour tissue dissociated at 37°C with collagenase or 6°C with cold active protease and stained with markers for tumour cells (EpCAM), endothelial cells (CD31), fibroblasts (FAP), lymphocytes (CD45), B Cells (CD19), NK cells (CD56) and T cells (CD8, CD3). **C** UMAP of combined scRNAseq experiments of ovarian cancer (n=2) and breast cancer (n-3) patient tissues with cell type assignments according to known gene markers for each cell type. **D** The top 40 genes from the geneset derived in Figure 3 as expressed in each cell type in breast and ovarian patient samples. Black circles around points denote significance at 5% FDR. **E** Pathway analysis of the differential expression results with the MSigDB hallmark gene sets for each cell type.

To uncover whether the transcriptional response to 37°C collagenase dissociation identified in PDX models is conserved in primary tumour samples, we next performed a differential expression analysis comparing the dissociation methods separately for each cell type (Figure 4 D). We found large consistent upregulation of the of the 512 genes identified in the core collagenase-associated geneset in PDX samples, with 61.7% to 78.1% upregulated across cell types and 8.6% to 54.9% significantly upregulated (table S4, Figures S7 and S8).

Though cell-type specific gene expression effects in response to digestion method were evident (Figure S9), global pathway analysis of differentially expressed genes for each cell type revealed conserved upregulation in NFκB signaling, apoptosis and inflammatory pathways as the most upregulated in all cell types (Figure 4 E). Smaller cell type-specific effects observed included increased hedgehog and apical surface pathways in breast epithelial cells, and reactive oxygen species pathways in cytotoxic T cells and myofibroblasts (Figure 4 E). Taken together, these findings indicate that all cell types exhibit some level of stress response to dissociation with collagenase, with some cell types exhibiting cell-type specific responses.

## Discussion

The advent of single cell sequencing technologies has empowered the study of complex biological systems including tissue microenvironments, tumour heterogeneity as well as the discovery of novel cell types otherwise difficult to detect [1]. Current sequencing techniques require single cell suspensions for passage through microfluidic or microwell platforms, and generation of single cell suspensions from solid tissues requires the enzymatic and mechanical disruption of extracellular matrix and cell-cell contacts. To date, the effect of these dissociation methods on the transcriptome of single cells has been largely ignored, despite the potential effects on the interpretation of scRNAseq data. Moreover, during both dissociation of tissues and passage through fluidic devices, cells can undergo stress, shearing, anoikis and apoptosis [26]. For this reason, efforts must be made on both sample handling and bioinformatics to ensure minimal noise and optimal filtration of data. Here, we endeavoured to describe the artifactual gene expression associated with tissue dissociation and dead or dying cell populations.

Using a large, diverse dataset, we highlight the variability in key QC metrics, including percentage of mitochondrial genes, number of UMIs and number of genes detected. We identify subpopulations of dead cells that express either high or low mitochondrial genes, contrary to the notion that dead cells can be characterized by their mitochondrial gene content alone. Importantly, cells that are FACS sorted as either live, dying, or dead based on PI/Annexin V staining are present in all three clusters, highlighting that the transcriptomic state of the cell is not necessarily the same as the surface marker state (though the two are correlated). As noted, this is reminiscent of “pseudotime” orderings, with PC1 from **Figure 2A** approximating a trajectory through the data that tracks transcriptomically healthy cells to transcriptomically dead cells with increasing PC1 values. Though transcriptomally similar to live, healthy cells, dead cells with low mitochondrial content expressed significantly high levels of MHC class I genes such as *HLA-A*, *HLA-B* and *B2M*. MHC class I genes are involved in antigen presentation to T cells, but are also expressed in many cell types and are induced in response to stress stimuli and contain heat shock-inducible elements [19]. In addition to standard practices of excluding cells with high mitochondrial content, cells with induction of these MHC class I genes may also be considered with caution. Moreover, interpretation of stress pathway expression in single cell studies, including components of surface immune recognition such as MHC class I, may be especially confounded.

We identify a conserved collagenase-associated transcriptional pattern including induction of stress and heat shock genes, consistent with a transcriptional response identified in a subset of muscle stem cells [23], and which was minimized when samples were dissociated at cold temperatures with a cold active serine protease. Transcription of these genes as a result of sample preparation methods may mask their induction due to other means. For example, *JUN* and *FOS* are associated with cancer drug resistance and metastatic progression [27–29]. Moreover, though less stark as the core collagenase-associated geneset, cell type-specific effects were observed during dissociation, included increased hedgehog and apical surface pathways in breast epithelial cells, and reactive oxygen species pathways in cytotoxic T cells and myofibroblasts. Taken together, these findings indicate that all cell types exhibit some level of stress response to dissociation with collagenase, with some cell types exhibiting cell-type specific responses. These stress responses, which may significantly influence the interpretation of scRNAseq data, are minimized by dissociation at cold temperatures.

## Methods

### Ethical approval

The Ethics Committees at the University of British Columbia approved all the experiments using human resources. Patients in Vancouver, British Columbia were recruited and samples were collected under tumour tissue repository (H06-00289) and Neoadjuvant PDX (H11-01887) protocol with informed consent. This fulfills the requirements of UBC BCCA Research Ethics Board. All animal studies were approved by the Animal Care Committee at the University of British Columbia.

### Specimen collection

After informed consent, tumour fragments from patients undergoing excision or diagnostic core biopsy were collected. Tumour materials were processed as described in [30].

### Patient-derived Xenografts

Tumour fragments were transplanted subcutaneously into female NOD/SCID interleukin-2 receptor gamma null (NSG) and NOD Rag-1 null interleukin-2 receptor gamma null (NRG) mice as previously described [30].

### Tissue dissociation at 37°C

Tumour fragments from patient breast and ovarian samples and PDXs were incubated for 2 hrs with a collagenase/hyaluronidase enzyme mix in serum-free Dulbecco’s Modified Eagle’s Medium (DMEM) at 37°C with intermittent gentle trituration with a wide bore pipette tip. Cells were resuspended in 0.25% trypsin-edta for 1 min followed by neutralization with 2% FBS in Hank’s Balanced Salt Solution (HBSS) and centrifugation. Cells were resuspended in 2% FBS/HBSS and filtered through a 40µm filter. Where necessary, dead cells were removed using MACS Dead Cell Removal Beads (Miltenyi Biotec) according to the manufacturer’s instructions. Cells were centrifuged and resuspended in 0.04% BSA/PBS and cell concentration adjusted for scRNAseq.

### Tissue dissociation at 6°C

Tumour fragments were incubated for 30 mins at 6°C with a serine protease, subtilisin A, derived from the Himalayan soil bacterium *Bacillus Lichenformis* (Creative Enzymes NATE0633) in PBS supplemented with 5mM CaCl2 and 125U/ml DNAse, as described in [6, 31]. During dissociation, samples were gently triturated every 5 min using a wide-bore pipette. Cells were resuspended in 0.25% trypsin-edta for 1 min at room temperature, neutralized with 2% FBS in HBSS and filtered through a 40µm filter. Following dissociation, samples were processed for scRNAseq as described above.

### Cell culture

GM18507 cells were maintained in RPMI-1640 supplemented with 10% FBS. MDA-MB-231 cells were maintained in DMEM supplemented with 10% FBS. Cells were trypsinized using 0.05% trypsin-edta and placed on ice. Cells were then incubated for 2 hrs at 6°C, 24°C, 37°C or 42°C before being harvested for scRNAseq.

### Flow Cytometry

GM18507 cells were treated with or without 100ng/ml TNFαfor 24 hrs before being stained with propidium iodide and annexin V and sorted into dying, dead or live populations according to single, double or negative staining respectively using a FACS Aria Fusion (BD Biosciences).

### Single cell RNA sequencing

Single cell suspensions were loaded onto a 10x Genomics Chromium single cell controller and libraries prepared according to the 10x Genomics Single Cell 3’ Reagent kit standard protocol. Libraries were then sequenced on an Illumina Nextseq500/550 with 42bp paired end reads, or a HiSeq2500 v4 with 125bp paired end reads. 10x Genomics Cell Ranger 3.0.2 was used to perform demultiplexing, counting and alignment to GRCh38 and mm10.

### Removal of murine contamination from patient derived xenograft samples

To identify murine cells in the PDX samples, we re-ran CellRanger version 3.0.2 aligning cells to both GRCh38 and mm10 (separately). We then considered all cells for which a valid barcode was identified in the raw (unfiltered) data for either alignment, and counted the number of reads mapping to each genome for each cell. A cell was subsequently designated as a contaminating mouse cell if more reads mapped to mm10 than GRCh38, and a human cell otherwise.

### Analysis of existing 10X datasets

The processed data for the datasets nuclei_900, pbmc4k, t_4 were downloaded from the 10X genomics website https://support.10xgenomics.com/single-cell-gene-expression/datasets/2.1.0/ on April 30th 2019.

### Differential expression and core heat-related gene set

All differential expression analyses were performed with edgeR [22] version 3.24.3 using the quasi-likelihood F-test as was the top-performing method in a recent review [32]. We included both a scaled cellular detection rate (scaled fraction of transcriptome detected per cell) and the patient / xenograft / cell line ID in the design matrix to account for unwanted technical and biological variation. In every case we only considered genes with minimum 10 counts across all cells.

We defined the core set of genes as those with FDR adjusted Q-value < 0.05 and with | log_2_(fold change)| > log 2(1.5) - in other words we require the average change in expression to be either 50% greater or less than the baseline to include the gene. Overall this gave 192 genes (182 upregulated and 10 downregulated).

Pathway enrichment was performed using camera [33] with trend.var=TRUE on the Hallmark gene set [24] retrieved from http://bioinf.wehi.edu.au/software/MSigDB/human_H_v5p2.rdata with timestamp 2016-10-10.

### Cell type assignments

Cell types were determined using CellAssign, a probabilistic model that annotates scRNAseq data into pre-defined and de novo cell types assuming a set of markers known marker genes for cell types [25]. Briefly, CellAssign takes a pre-defined set of marker genes for each cell type in the data, and probabilistically models a cell as being of a certain type if it has increased expression of its marker genes. A given gene can be a marker for multiple cell types and a marker gene can be expressed in cell types other than those for which it is a marker, albeit at lower levels. The marker genes used in this study are listed in table S2 and table S3.

### Clustering of live, dying, and dead cells

Cells were hierarchically clustered using the `hclust` function in `R` applied to the 10-dimensional output of MNN, and clusters assigned using the `cutree` function.

### Reproducible data analysis

A dockerized workflow to enable reproduction of all figures and analysis in this paper is available at https://github.com/kieranrcampbell/scrnaseq-digestion-paper with corresponding docker image at https://cloud.docker.com/u/kieranrcampbell/repository/docker/kieranrcampbell/statgen2 (version 0.4).

## Supporting information

Core collagenase 37C gene set

## Abbreviations

UMI: unique molecular identifier
PDX: patient-derived xenograft
MHC: major histo-compatibility complex
QC: quality control
PCA: principal componenst analysis
HSP: heat shock protein
UMAP: uniform manifold approximation and projection
FDR: false discovery rate
HGSC: high grade serous carcinoma

## Acknowledgements

This work was supported by the BC Cancer Foundation, Canadian Institutes for Health Research (CIHR), Canadian Cancer Society Research Institute (CCSRI), Terry Fox Research Institute (TFRI), Canadian Foundation for Innovation (CFI), Canada Research Chairs program, Michael Smith Foundation for Health Research (MSFHR) and Cancer Research UK (Grand challenge IMAXT award). KRC is funded by postdoctoral fellowships from the Canadian Institutes of Health Research, the Canadian Statistical Sciences Institute (CANSSI), and the UBC Data Science Institute. SPS is a Susan G. Komen scholar.

## Authors’ contributions

SA and SPS conceived of and oversaw the study. COF, FK, JL, JB and PE conducted the experiments. COF conducted the single cell experiments. FK generated the PDX tissues. KC and AWZ analyzed the data. RM and AJM performed the sequencing. DL, AM, MW, JH, BH, NC performed the data processing. EK, KB, CB and JM provided the primary tumour tissue. All authors read and approved the final manuscript.

## Ethical approval

The Ethics Committees at the University of British Columbia approved all the experiments using human resources.

## Consent for publication

Not applicable

## Availability of data and material

**Figure S1:**
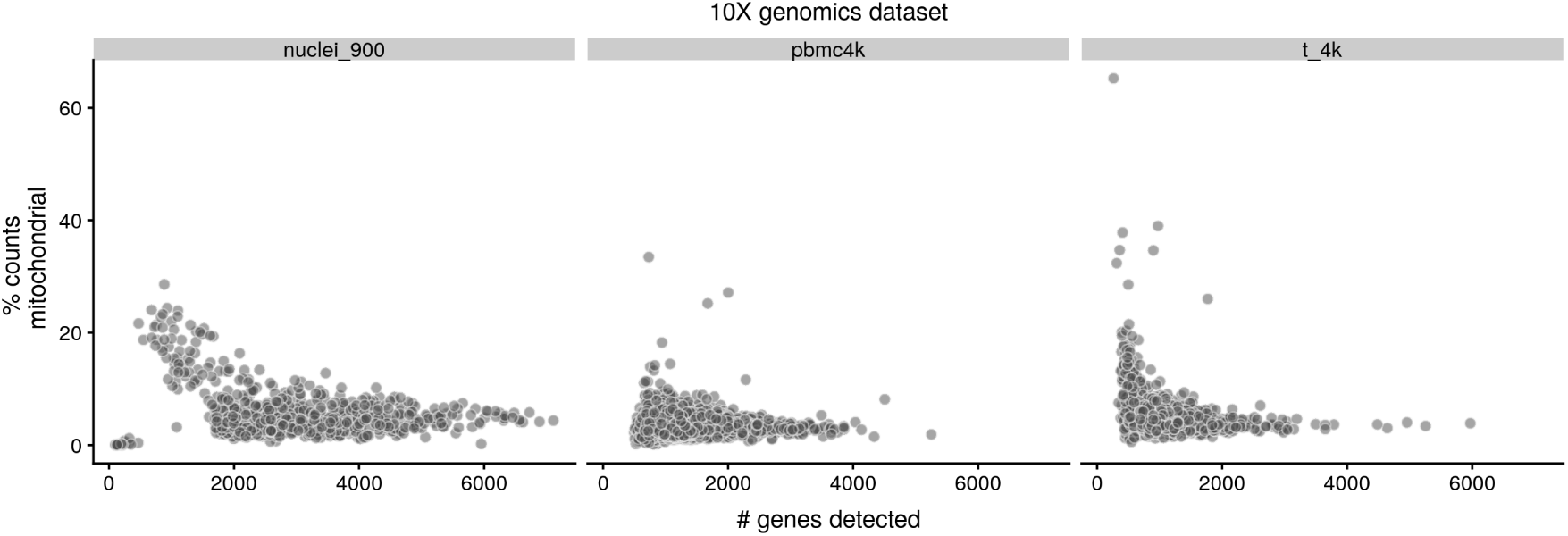
Mitochondrial gene content of single cells as a function of number of genes detected for three publicly available datasets using the 10X genomics platform (nuclei_900, pbmc4k, t_4). These datasets contain sets of cells with distinctly increased mitochondrial gene content percentages with a lower number of detected genes.

**Figure S2:**
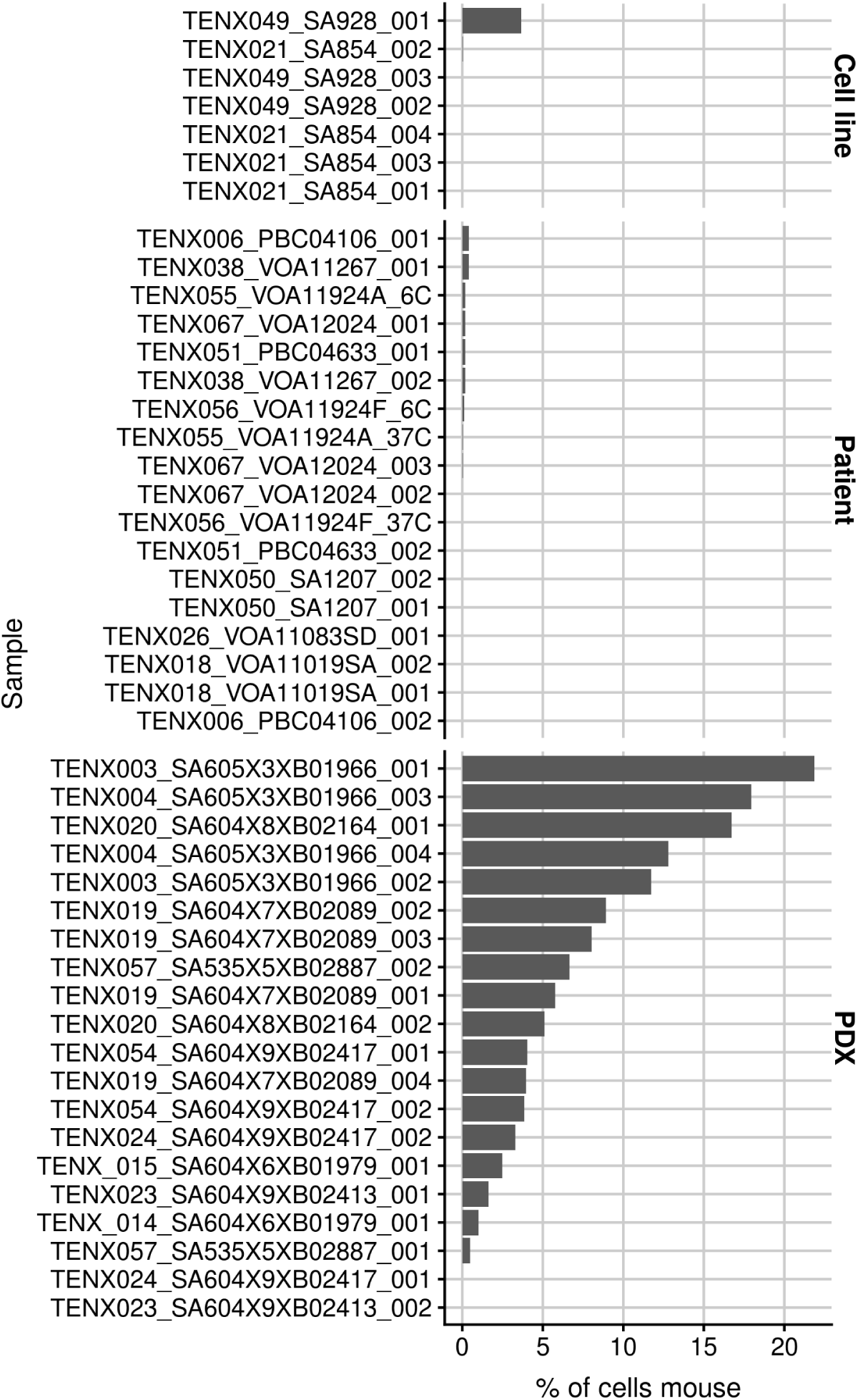
Percentage of cells identified as mouse cells across all scRNA-seq samples in the study exhibits large variation across datasets. While the vast majority were detected in PDX samples, a small number were also detected in cell line and patient primary tumour samples, suggesting either small levels of contamination or a modest false positive rate to the detection method.

**Figure S3:**
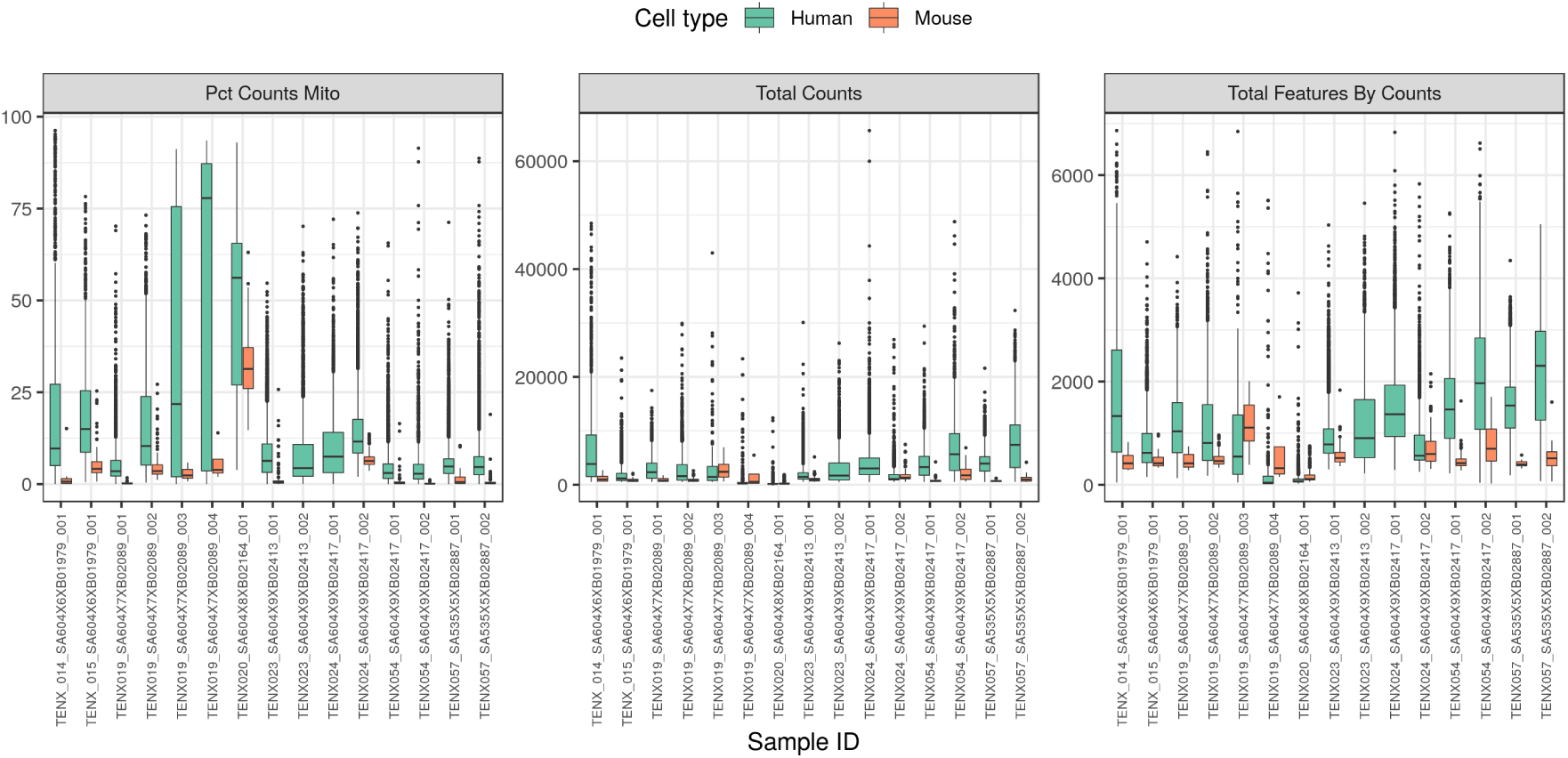
Quality control metrics (% mitochondrial gene counts, total counts, total genes detected) across all scRNA-seq of PDX included in the study. On average, murine cells score lower across all three metrics though with notable inter-dataset variability.

**Figure S4:**
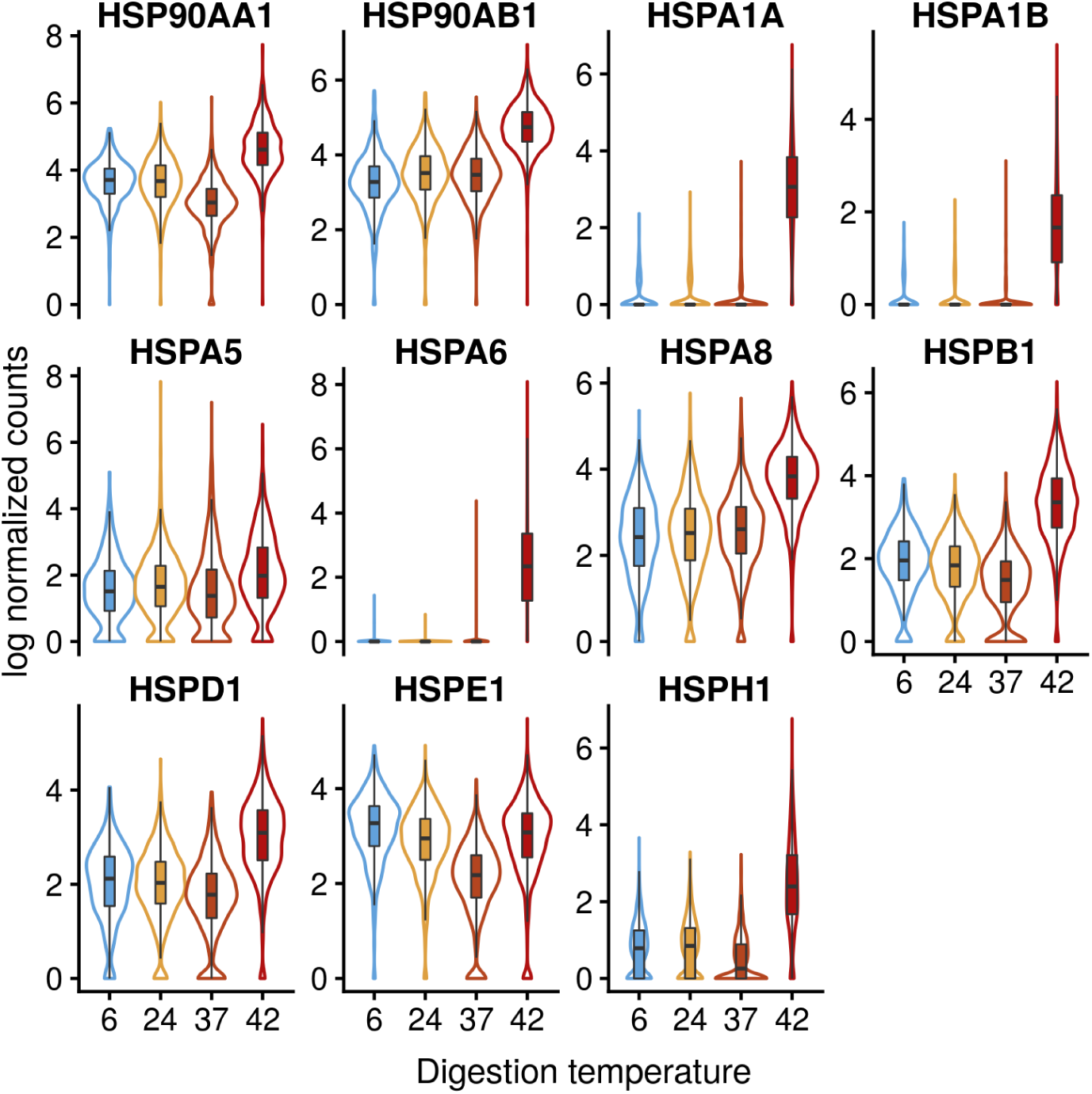
Heat shock protein (HSP) gene expression across different digestion temperatures in MDA-MB-231 cells.

**Figure S5:**
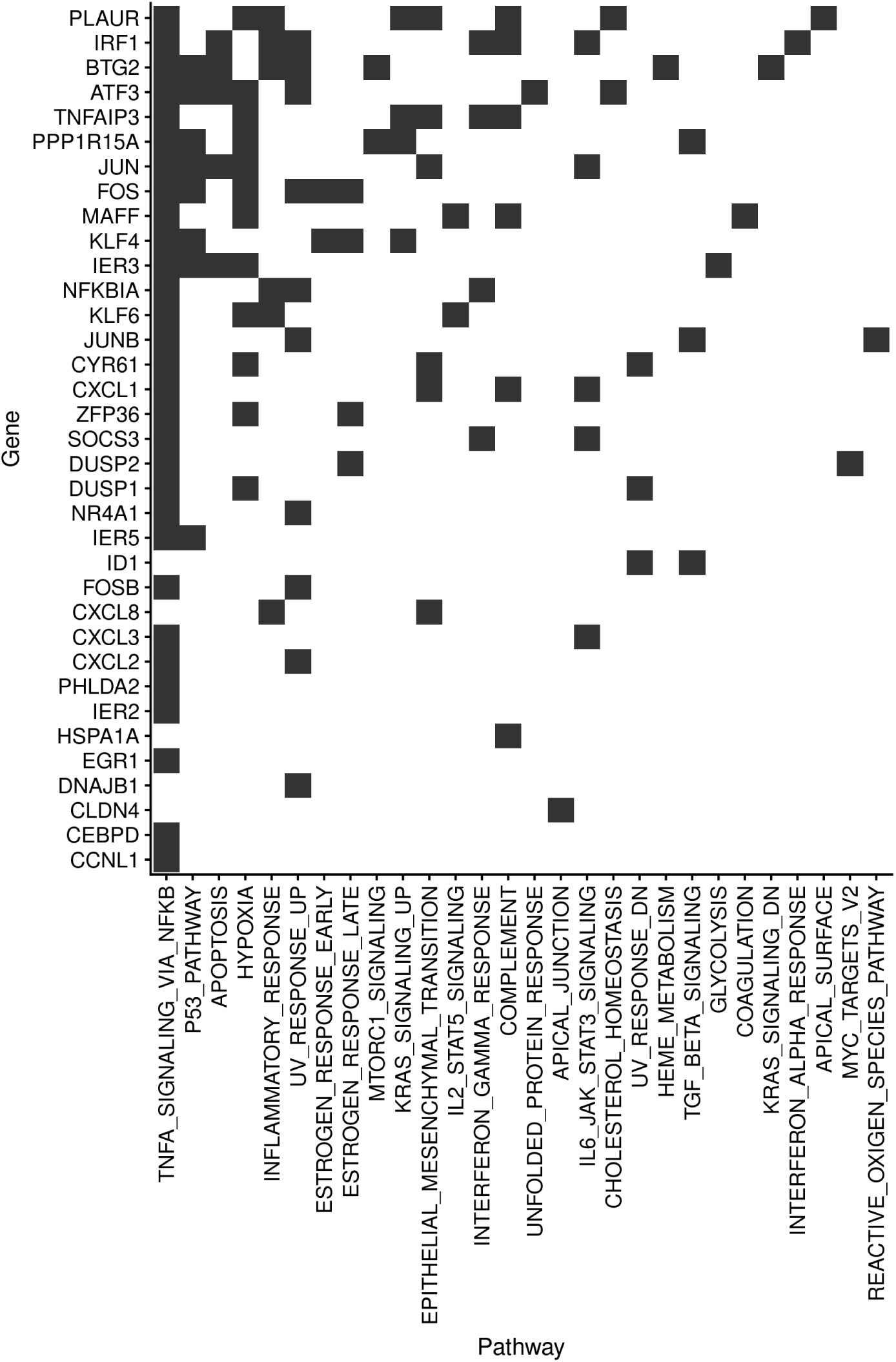
Pathway membership of top 40 genes (based on log fold change) from the 512 genes in the PDX digestion method core set.

**Figure S6:**
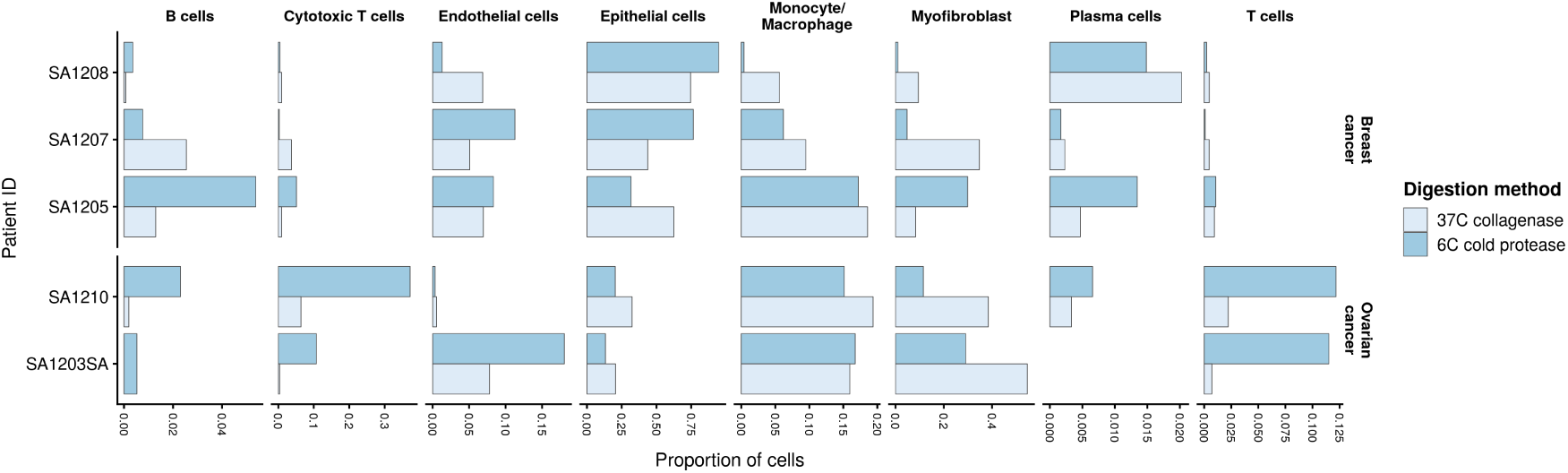
Microenvironment composition across primary tumour samples digested at 6 °C or 37 °C. Results show enhanced capture of T cell and cytotoxic T cells in ovarian cancer patient samples, though the composition of the tumour samples was variable, and no consistent difference was observed between conditions in all samples.

**Figure S7:**
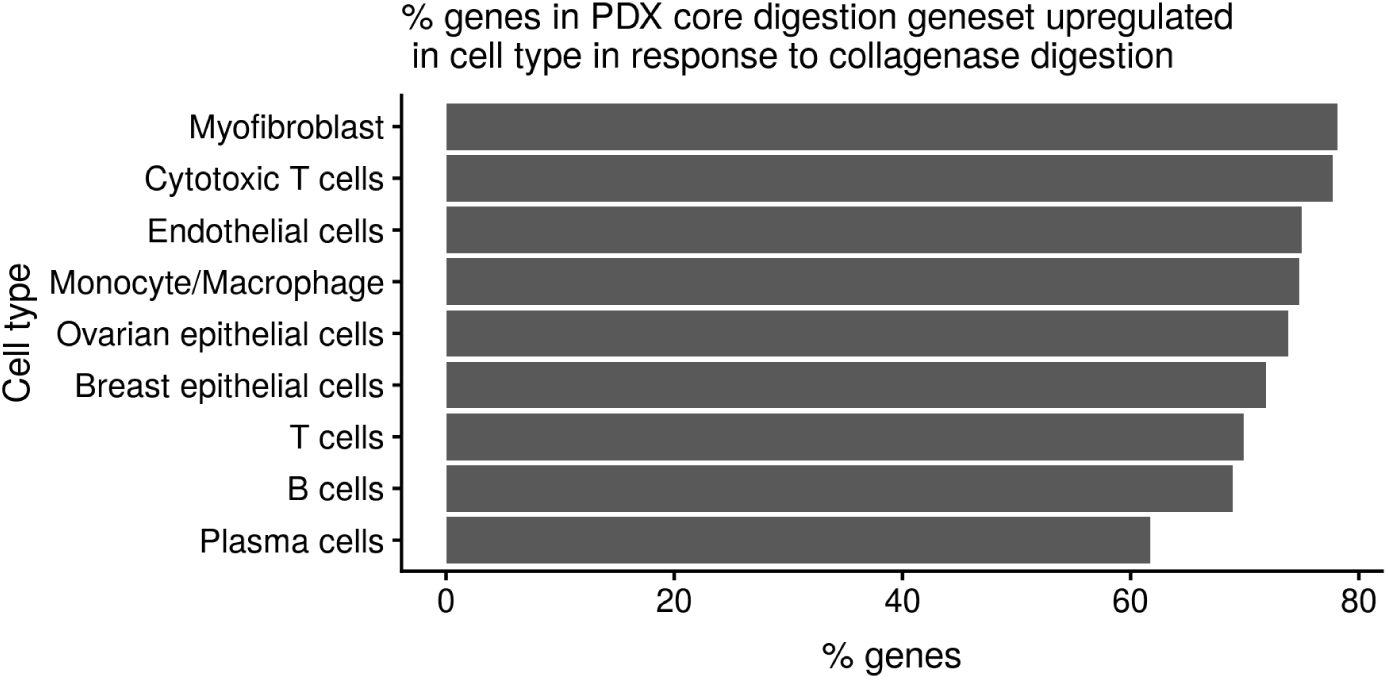
% of genes identified in the 512 digestion method dependent core gene set in PDX that are upregulated in each primary tumour cell type.

**Figure S8:**
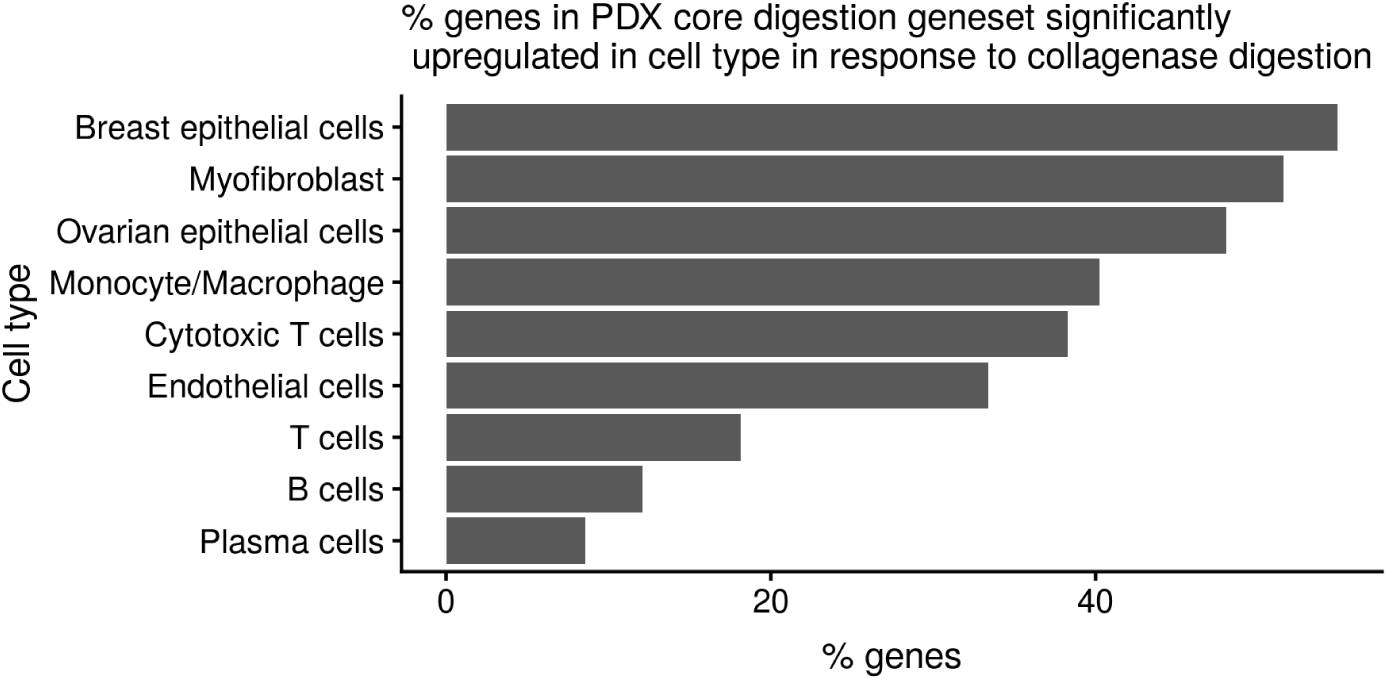
% of genes identified in the 512 digestion method dependent core gene set in PDX that are significanty upregulated (FDR < 5%) in each primary tumour cell type.

**Figure S9:**
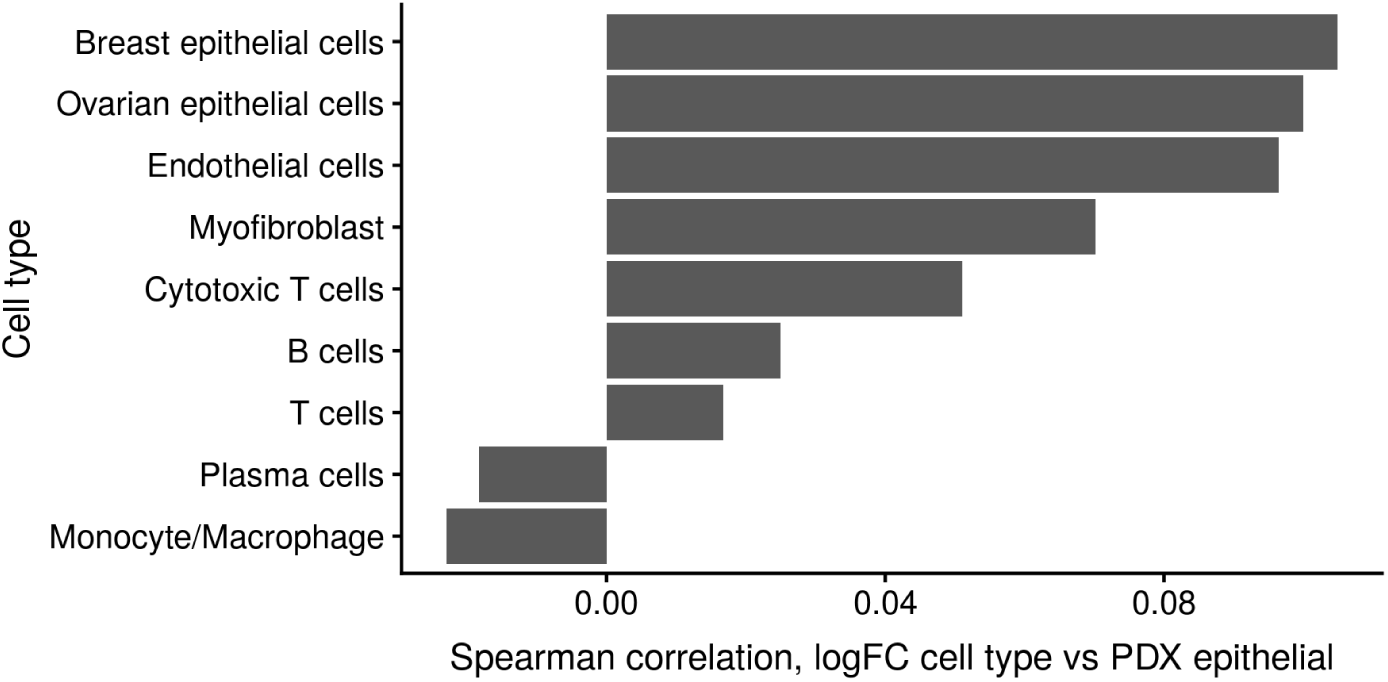
Spearman correlation between the log fold changes in response to digestion method (37C collagenase vs 6C cold protease) in PDX vs. primary tumours of a given cell type.

**Table S1:**
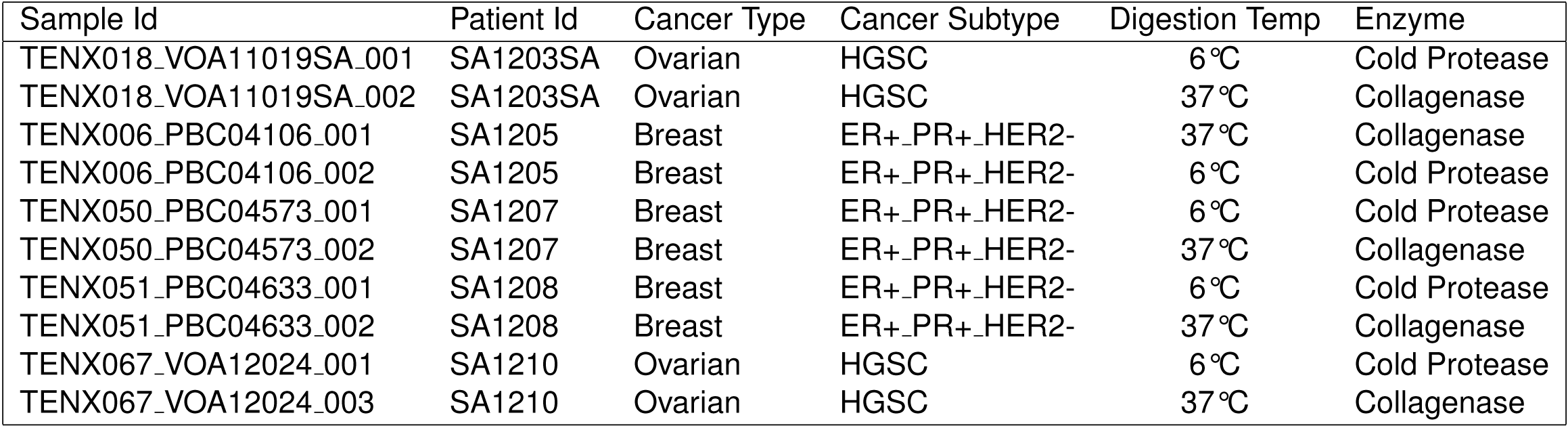
Characteristics of primary tumour samples

**Table S2:** Marker genes used for cell type assignments of breast patient samples

| Cell Type | Markers |
| --- | --- |
| B cell | VIM, MS4A1, CD79A, PTPRC, CD19, BANK1, IGKC, LAPTM5 |
| Plasma cell | VIM, CD79A, PTPRC, SDC1, IGKC, IGHG1, IGHG2, LAPTM5 |
| T cells | VIM, CD2, CD3D, CD3E, CD3G, CD28, CD4, PTPRC, LAPTM5 |
| Cytotoxic T cells | VIM, CD2, CD3D, CD3E, CD3G, CD28, CD8A, GZMA, PRF1, NKG7, PTPRC, GZMB, LAPTM5 |
| Monocyte/Macrophage | VIM, CD14, FCGR3A, CD33, ITGAX, ITGAM, CD4, PTPRC, LYZ, LAPTM5 |
| Breast cancer cells | EPCAM, KRT18, KRT8, KRT19, ESR1 |
| Myofibroblast | VIM, ACTA2, COL1A1, COL3A1, SERPINH1, SDC1 |
| Endothelial cells | VIM, EMCN, CLEC14A, CDH5, PECAM1, VWF, SERPINH1, IL3RA |

**Table S3:** Marker genes used for cell type assignments of breast patient samples

| Cell Type | Markers |
| --- | --- |
| B cells | VIM, MS4A1, CD79A, PTPRC, CD19, BANK1, CD24, IGKC |
| Plasma cells | VIM, CD79A, PTPRC, SDC1, IGKC, IGHG1, IGHG2 |
| T cells | VIM, CD2, CD3D, CD3E, CD3G, CD28, CD4, PTPRC |
| Cytotoxic T cells | VIM, CD2, CD3D, CD3E, CD3G, CD28, CD8A, GZMA, PRF1, NKG7, PTPRC, GZMB |
| Monocyte/Macrophage | VIM, CD14, FCGR3A, CD33, ITGAX, ITGAM, CD4, PTPRC, LYZ |
| Epithelial cells | EPCAM, WFDC2, CD24, CDH1, CLDN3, CLDN4 |
| Myofibroblast | VIM, ACTA2, COL1A1, COL3A1, SERPINH1, SDC1 |
| Endothelial cells | VIM, EMCN, CLEC14A, CDH5, PECAM1, VWF, SERPINH1, IL3RA |

**Table S4:** % of the 512 genes in the 37°C, collagenase-associated digestion core gene set identified in PDX samples upregulated and significantly upregulated (5% FDR) in each primary tumour cell type.

| Cell type | % upregulated | % significantly upregulated |
| --- | --- | --- |
| B cells | 68.95 | 12.11 |
| Breast epithelial cells | 71.88 | 54.88 |
| Cytotoxic T cells | 77.73 | 38.28 |
| Endothelial cells | 75.00 | 33.40 |
| Monocyte/Macrophage | 74.80 | 40.23 |
| Myofibroblast | 78.12 | 51.56 |
| Ovarian epithelial cells | 73.83 | 48.05 |
| Plasma cells | 61.72 | 8.59 |
| T cells | 69.92 | 18.16 |

## References

1. S Steven Potter. “Single-cell RNA sequencing for the study of development, physiology and disease”. In: Nature Reviews Nephrology (2018), p. 1.

2. Oliver Stegle, Sarah A Teichmann, and John C Marioni. “Computational and analytical challenges in single-cell transcriptomics”. In: Nature Reviews Genetics 16.3 (2015), p. 133.

3. Arjun Raj and Alexander van Oudenaarden. “Nature, nurture, or chance: stochastic gene expression and its consequences”. In: Cell 135.2 (2008), pp. 216–226.

4. Saiful Islam, et al. “Quantitative single-cell RNA-seq with unique molecular identifiers”. In: Nature methods 11.2 (2014), p. 163.

5. Ilan Volovitz, et al. “A non-aggressive, highly efficient, enzymatic method for dissociation of human brain-tumors and brain-tissues to viable single-cells”. In: BMC neuroscience 17.1 (2016), p. 30.

6. Mike Adam, Andrew S Potter, and S Steven Potter. “Psychrophilic proteases dramatically reduce single cell RNA-seq artifacts: A molecular atlas of kidney development”. In: Development (2017), dev–151142.

7. Sohrab P Shah, et al. “The clonal and mutational evolution spectrum of primary triple-negative breast cancers”. In: Nature 486.7403 (2012), p. 395.

8. Danny R Welch. “Tumor heterogeneity—a ‘contemporary concept’founded on historical insights and predictions”. In: Cancer research 76.1 (2016), pp. 4–6.

9. Devon A Lawson, et al. “Tumour heterogeneity and metastasis at single-cell resolution”. In: Nature cell biology 20.12 (2018), p. 1349.

10. Elham Azizi, et al. “Single-cell map of diverse immune phenotypes in the breast tumor microenvironment”. In: Cell 174.5 (2018), pp. 1293–1308.

11. Charissa Kim, et al. “Chemoresistance evolution in triple-negative breast cancer delineated by single-cell sequencing”. In: Cell 173.4 (2018), pp. 879–893.

12. Grace XY Zheng, et al. “Massively parallel digital transcriptional profiling of single cells”. In: Nature communications 8 (2017), p. 14049.

13. Leland McInnes and John Healy. “Umap: Uniform manifold approximation and projection for dimension reduction”. In: arXiv preprint arXiv:1802.03426 (2018).

14. Tomislav Ilicic, et al. “Classification of low quality cells from single-cell RNA-seq data”. In: Genome biology 17.1 (2016), p. 29.

15. Quan Zhao et al. “A mitochondrial specific stress response in mammalian cells”. In: The EMBO Journal 21.17 (2002), pp. 4411–4419. ISSN: 0261-4189. DOI: 10.1093/emboj/cdf445. eprint: http://emboj.embopress.org/content/21/17/4411.full.pdf. URL: http://emboj.embopress.org/content/21/17/4411.

16. EA Carswell, et al. “An endotoxin-induced serum factor that causes necrosis of tumors”. In: Proceedings of the National Academy of Sciences 72.9 (1975), pp. 3666–3670.

17. Lisa M Sedger and Michael F McDermott. “TNF and TNF-receptors: from mediators of cell death and inflammation to therapeutic giants–past, present and future”. In: Cytokine & growth factor reviews 25.4 (2014), pp. 453–472.

18. Laleh Haghverdi, et al. “Batch effects in single-cell RNA-sequencing data are corrected by matching mutual nearest neighbors”. In: Nature biotechnology 36.5 (2018), p. 421.

19. Michael Gleimer and Peter Parham. “Stress management: MHC class I and class I-like molecules as reporters of cellular stress”. In: Immunity 19.4 (2003), pp. 469–477.

20. Kieran R Campbell and Christopher Yau. “A descriptive marker gene approach to single-cell pseudotime inference”. In: Bioinformatics 35.1 (2018), pp. 28–35.

21. Kieran R Campbell and Christopher Yau. “Uncovering pseudotemporal trajectories with covariates from single cell and bulk expression data”. In: Nature communications 9.1 (2018), p. 2442.

22. Mark D Robinson, Davis J McCarthy, and Gordon K Smyth. “edgeR: a Bioconductor package for differential expression analysis of digital gene expression data”. In: Bioinformatics 26.1 (2010), pp. 139–140.

23. Susanne C van den Brink et al. “Single-cell sequencing reveals dissociation-induced gene expression in tissue subpopulations”. In: Nature methods 14.10 (2017), p. 935.

24. Arthur Liberzon, et al. “The molecular signatures database hallmark gene set collection”. In: Cell systems 1.6 (2015), pp. 417–425.

25. Allen W Zhang, et al. “Probabilistic cell type assignment of single-cell transcriptomic data reveals spatiotemporal microenvironment dynamics in human cancers”. In: bioRxiv (2019), p. 521914.

26. Aisha A AlJanahi, Mark Danielsen, and Cynthia E Dunbar. “An Introduction to the analysis of single-cell RNA-sequencing data”. In: Molecular Therapy-Methods & Clinical Development 10 (2018), pp. 189–196.

27. Jacob Insua-Rodrıguez, et al. “Stress signaling in breast cancer cells induces matrix components that promote chemoresistant metastasis”. In: EMBO molecular medicine 10.10 (2018), e9003.

28. F Fan, et al. “The AP-1 transcription factor JunB is essential for multiple myeloma cell proliferation and drug resistance in the bone marrow microenvironment”. In: Leukemia 31.7 (2017), p. 1570.

29. Rachel Ramsdale, et al. “The transcription cofactor c-JUN mediates phenotype switching and BRAF inhibitor resistance in melanoma”. In: Sci. Signal. 8.390 (2015), ra82–ra82.

30. Peter Eirew, et al. “Dynamics of genomic clones in breast cancer patient xenografts at single-cell resolution”. In: Nature 518.7539 (2015), p. 422.

31. Andrew S Potter and S Steven Potter. “Dissociation of Tissues for Single-Cell Analysis”. In: Kidney Organogenesis. Springer, 2019, pp. 55–62.

32. Charlotte Soneson and Mark D Robinson. “Bias, robustness and scalability in single-cell differential expression analysis”. In: Nature methods 15.4 (2018), p. 255.

33. Di Wu and Gordon K Smyth. “Camera: a competitive gene set test accounting for inter-gene correlation”. In: Nucleic acids research 40.17 (2012), e133–e133.

